# Compendiums of Cancer Transcriptome for Machine Learning Applications

**DOI:** 10.1101/353698

**Authors:** Su Bin Lim, Swee Jin Tan, Wan-Teck Lim, Chwee Teck Lim

**Affiliations:** NUS Graduate School for Integrative Sciences & Engineering, National University of Singapore, #05-01, 28 Medical Drive, Singapore, 117456 Singapore; Department of Biomedical Engineering, National University of Singapore, 4 Engineering Drive 3, Engineering Block 4, #04-08, Singapore, 117583 Singapore; Sysmex Asia Pacific Pte Ltd, 9 Tampines Grande, #06-18, Singapore, 528735 Singapore; Division of Medical Oncology, National Cancer Centre Singapore, 11 Hospital Drive, Singapore, 169610 Singapore; Office of Clinical Sciences, Duke-NUS Medical School, 8 College Road, Singapore, 169857 Singapore; Institute of Molecular and Cell Biology, A*Star, 61 Biopolis Drive, Proteos, Singapore, 138673 Singapore; Mechanobiology Institute, National University of Singapore, #10-01, 5A Engineering Drive 1, Singapore, 117411 Singapore; Biomedical Institute for Global Health Research and Technology, National University of Singapore, #14-01, MD6, 14 Medical Drive, Singapore, 117599 Singapore

**Keywords:** Digital curation, data integration, machine learning, cross-platform analysis, cancer

## Abstract

**Background:** There exist massive transcriptome profiles in the form of microarray, enabling reuse. The challenge is that they are processed with diverse platforms and preprocessing tools, requiring considerable time and informatics expertise for cross-dataset or cross-cancer analyses. If there exists a single, integrated data source consisting of thousands of samples, similar to TCGA, data-reuse will be facilitated for discovery, analysis, and validation of biomarker-based clinical strategy.

**Findings:** We present 11 merged microarray-acquired datasets (MMDs) of major cancer types, curating 8,386 patient-derived tumor and tumor-free samples from 95 GEO datasets. Highly concordant MMD-derived patterns of genome-wide differential gene expression were observed with matching TCGA cohorts. Using machine learning algorithms, we show that clinical models trained from all MMDs, except breast MMD, can be directly applied to RNA-seq-acquired TCGA data with an average accuracy of 0.96 in classifying cancer. Machine learning optimized MMD further aids to reveal immune landscape of human cancers critically needed in disease management and clinical interventions.

**Conclusions:** To facilitate large-scale meta-analysis, we generated a newly curated, unified, large-scale MMD across 11 cancer types. Besides TCGA, this single data source may serve as an excellent training or test set to apply, develop, and refine machine learning algorithms that can be tapped to better define genomic landscape of human cancers.

## Introduction

Although there exist vast datasets deposited at NCBI GEO in the form of microarray, applying machine learning to exploit them is not straightforward. They are often generated with diverse platforms and normalization tools, limited in sample size, and annotated with non-standardized text – all these add computational complexity to such high-dimensional data, necessitating multiple, intricate analytics tools at different steps for data integration.

To increase the reuse of such legacy data, we generated a single, merged microarray-acquired datasets (MMDs) of 11 major cancer types within a uniform R framework (Figure 1). This approach has been used in our earlier work to generate merged transcriptome data of specific cancer type, non-small cell lung cancer (NSCLC), comprising both tumor and tumor-free tissues [1]. Utilising this merged dataset, we identified a specific matrisome expression pattern predictive of prognosis and adjuvant chemotherapy response [1]. Such large-scale data allow detection of signals from genes that may be masked in smaller patient cohort.

**Figure 1.**
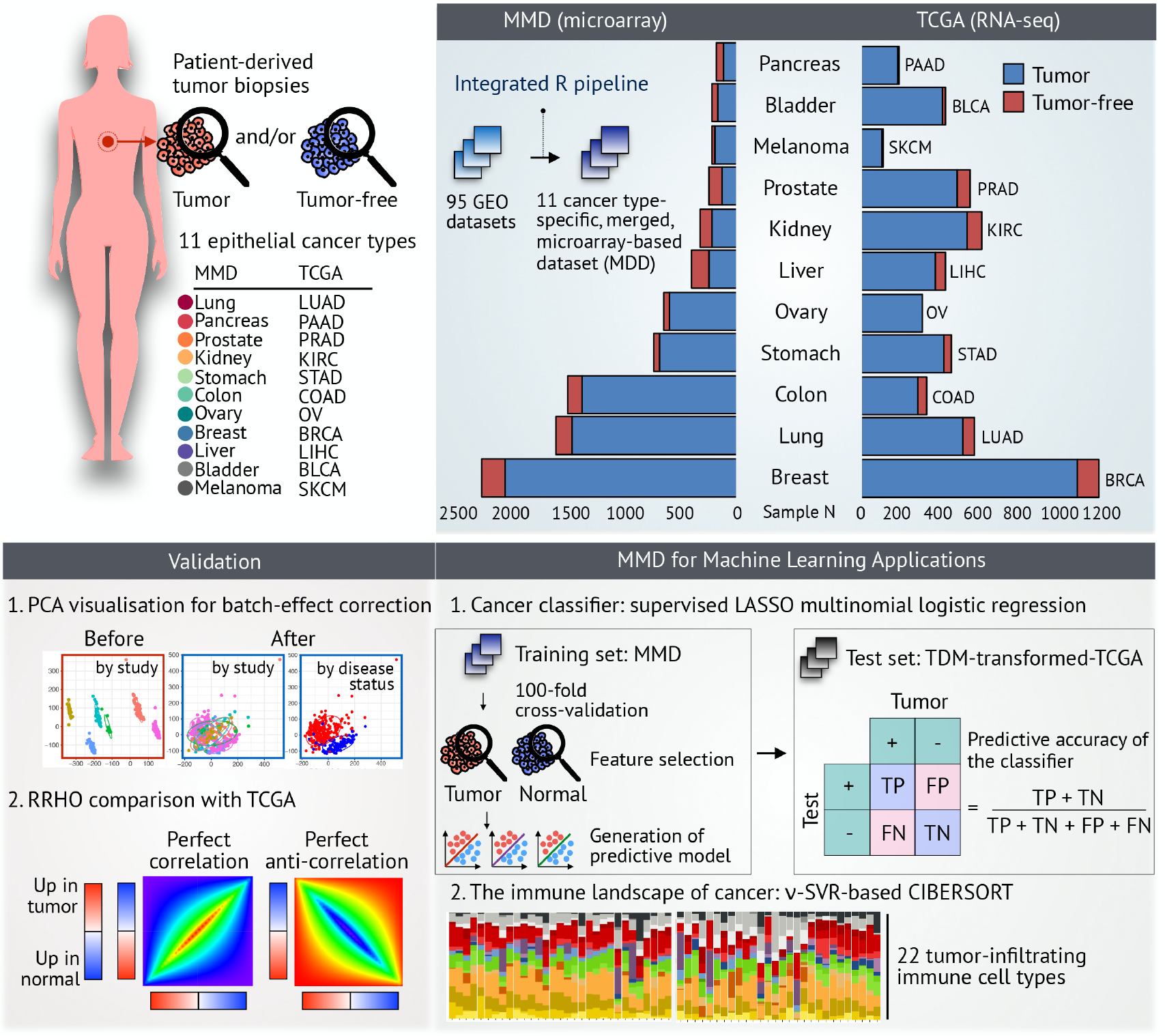
MMD: development, validation, and potential applications in oncology. Microarray-based datasets containing raw transcriptome profiles of patient-derived tumor and tumor-free tissues were processed, merged, and batch-effect corrected using an integrated R pipeline. Validation of each cancer type-specific MMD was performed using PCA and RRHO algorithms. Using machine learning, clinical models built from MMD can be applied to TCGA, allowing discovery of new biomarkers, development of prognostic model, and parallel cross-platform analyses with TCGA.

As an exemplary use of big data, TCGA increasingly serves as a ‘training’ reference to apply machine learning algorithms, having comprehensive, well-curated genomic data of over 11,000 tumors across 33 major cancer types. This rich resource combined with machine learning has facilitated recent development and/or validation of cancer type classifier [2], markers predictive of drug sensitivity [3], histopathology image-based prognostic predictor [4], and previously unexplored biological feature associated with oncogenic phenotype [5].

In this work we extend the algorithm to include various carcinomas of epithelial origin. Meeting our inclusion criteria, raw expression profiles of 8,386 patient-derived tumor and normal tissues obtained from 95 independent GEO datasets have been curated, merged, and batch-effect corrected. Consistent with prior works [6–10], we observed comparably correlated patterns of genome-wide differential expression between microarray and RNA-seq. With an appropriate machine learning and normalization technique, it is thus expected that MMD can be used to generate clinical predictors that can be applied cross platform, given sufficient overlap in the data distribution [11].

To demonstrate, we developed an integrated R pipeline to train and build clinical predictive models from MMDs and test with respective transformed TCGA datasets. Applying an independent machine learning optimization [12], we further show how MMD can be used to de-convolve tumor immune microenvironment by parsing specific subpopulations of infiltrating immune cells in a given bulk tumor sample. We applied CIBERSORT method and compared the distribution of estimated composition and functional activation of 22 immune cell types with that of matching TCGA cohorts. All 11 cancer type-specific MMDs with associated clinical metadata are available at *ArrayExpress* (see Availability of supporting data). Our resource of curated large-scale data and integrated machine learning approach may allow extraction of unbiased and clinically significant information, enabling precision medicine.

## Data Description

### Data Collection and Preprocessing

To facilitate the use of MMDs for machine learning applications and parallel crossplatform analyses with TCGA, we chose eleven major epithelial cancer types, in which matching datasets, consisting of tumor and/or tumor-free tissues, are available: lung (LUAD), pancreas (PAAD), prostate (PRAD), renal (KIRC), gastric (STAD), colorectal (COAD), ovarian (OV), breast (BRCA), liver (LIHC), bladder (BLCA), and melanoma (SKCM) cancer. Only datasets having raw expression data assayed with the GPL570 platform (Affymetrix Human Genome U133 Plus 2.0 Array) were considered for data integration. Altogether we identified 95 independent GEO datasets (http://www.ncbi.nlm.nih.gov/geo) to generate 11 cancer type-specific MMDs (Table 1).

**Table 1.** Summary of 11 MMDs and 11 TCGA cohorts: dataset accession code and number of samples processed to generate MMD.

| MMD | Breast | Lung | Colon | Stomach | Ovary | Liver | Kidney | Prostate | Melanoma | Bladder | Pancreas |
| --- | --- | --- | --- | --- | --- | --- | --- | --- | --- | --- | --- |
| GEO accession codes | GSE3744 | GSE10245 | GSE4107 | GSE15459 | GSE9899 | GSE29721 | GSE36895 | GSE17951 | GSE19234 | GSE31189 | GSE16515 |
|  | GSE19615 | GSE10445 | GSE4183 | GSE34942 | GSE10971 | GSE6222 | GSE23629 | GSE55945 | GSE23376 | GSE7476 | GSE15471 |
|  | GSE21653 | GSE10799 | GSE9348 | GSE62254 | GSE14001 | GSE40873 | GSE11151 | GSE3325 | GSE7553 | GSE31684 | GSE32676 |
|  | GSE12276 | GSE12667 | GSE13067 | GSE13911 | GSE14407 | GSE75271 | GSE53757 | GSE26910 | GSE15605 | GSE30522 | GSE22780 |
|  | GSE26639 | GSE18842 | GSE13294 | GSE35809 | GSE18520 | GSE41804 | GSE12606 | GSE45016 |  |  |  |
|  | GSE9195 | GSE19188 | GSE14333 | GSE19826 | GSE19352 | GSE62232 | GSE12090 | GSE17906 |  |  |  |
|  | GSE23177 | GSE28571 | GSE15960 | GSE38749 | GSE15578 | GSE45436 |  |  |  |  |  |
|  | GSE23593 | GSE31210 | GSE17536 |  | GSE12172 |  |  |  |  |  |  |
|  | GSE16446 | GSE33356 | GSE17537 |  | GSE19829 |  |  |  |  |  |  |
|  | GSE20685 | GSE50081 | GSE18105 |  | GSE30161 |  |  |  |  |  |  |
|  | GSE20711 | GSE37745 | GSE18088 |  |  |  |  |  |  |  |  |
|  | GSE7904 | GSE30219 | GSE20916 |  |  |  |  |  |  |  |  |
|  | GSE18864 |  | GSE23878 |  |  |  |  |  |  |  |  |
|  | GSE42568 |  | GSE33113 |  |  |  |  |  |  |  |  |
|  | GSE50567 |  | GSE26682 |  |  |  |  |  |  |  |  |
|  | GSE28821 |  |  |  |  |  |  |  |  |  |  |
|  | GSE5764 |  |  |  |  |  |  |  |  |  |  |
|  | GSE10780 |  |  |  |  |  |  |  |  |  |  |
|  | GSE22513 |  |  |  |  |  |  |  |  |  |  |
|  | GSE28796 |  |  |  |  |  |  |  |  |  |  |
| Total N of samples | 2302 | 1621 | 1514 | 737 | 647 | 401 | 323 | 237 | 214 | 212 | 178 |
| Tumor tissue N | 2088 | 1474 | 1393 | 691 | 593 | 244 | 219 | 121 | 194 | 161 | 108 |
| Tumor-free tissue N | 214 | 147 | 121 | 46 | 54 | 157 | 104 | 116 | 20 | 51 | 70 |
| TCGA | BRCA | LUAD | COAD | STAD | OV | LIHC | KIRC | PRAD | SKCM | BLCA | PAAD |
| Total N of samples | 1207 | 574 | 327 | 450 | 304 | 421 | 606 | 549 | 103 | 427 | 182 |
| Tumor tissue N | 1094 | 515 | 286 | 415 | 304 | 371 | 533 | 482 | 102 | 408 | 178 |
| Tumor-free tissue N | 113 | 59 | 41 | 35 | 0 | 50 | 73 | 67 | 1 | 19 | 4 |

A total of 8,386 samples assayed was imported into R Bioconductor [13] and RMA-normalized using the *affy* package [14]. Depending on cancer type, these pre-processed data were merged and ComBat-adjusted for batch-effect removal using the *inSilicoMerging* package [15]. Probes having maximum expression values were then collapsed to the genes for subsequent downstream analyses. To compare RNA-Seq and microarray in transcriptome profiling, we performed global differential expression (DE) analyses using the *lmFit* and *eBayes* functions in the *limma* package [16].

The Cancer Genome Atlas (TCGA) data were retrieved and processed via the *TCGA-Assembler* package [17]. All sample data having primary tumor and/or normal tissues were downloaded across 11 epithelial cancers for whole-transcriptome profiles together with clinical metadata (Table 1). Normalized RPKM count values were extracted using the *ProcessRNASeqData* function via the *TCGA-Assembler* package [17]. Only genes with at least 1 count per million (cpm) or RPMK value in at least 20% of total number of samples in each cohort were kept via the *edgeR* package [18]. Selected genes were normalized by Trimmed Mean of M-values (TMM) and subjected to DE analyses using the *voom* and *lmFit* functions in the *limma* package [16]. Of note, ovarian (OV) and melanoma (SKCM) TCGA cohorts were excluded in RRHO analyses due to lack of normal samples (Table 1). Disease status (cancer vs. normal) corresponding to the 11 epithelial organs were obtained with *DownloadBiospecimenClinicalData* function in the *TCGA-Assembler* package [17].

## Technical Validation of MMD

### 1. Visual Validation of Batch-Effect Removal

Processing techniques were assessed to visualize the batch-effect correction with principal component analysis (PCA) plots, as previously described [1]. The first two PCs that capture the most variance are shown for both untransformed and ComBat-transformed datasets (Figure 2). In each case, there is an apparent overlay of the variance between independent studies, which are rather separated by the disease status. PCA was performed using the *prcomp* function in the built-in R *stats* package and visualized using the *ggbiplot* package [19].

**Figure 2.**
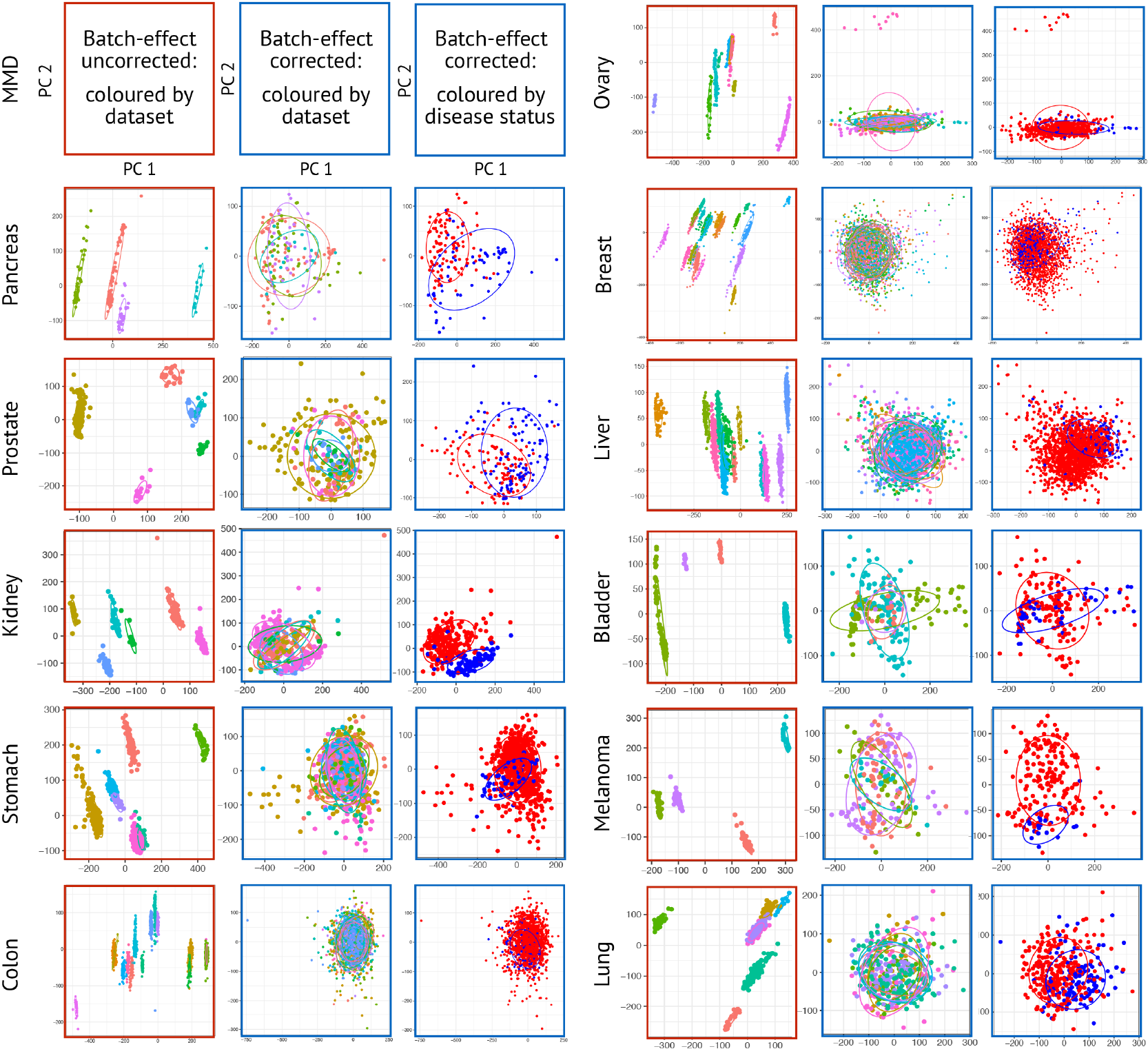
Visual validation of batch-effect removal. PCA plots showing the first two PCs which capture the most variance are shown. Plots with red colored-border show PCs before batch-effect correction and coloured by dataset. Plots with blue colored-border show PCs of ComBat-adjusted MMD and coloured by dataset and disease status (cancer vs. normal). Ellipses are drawn one standard deviation away from the mean of the Gaussian fitted to each independent MMD.

### 2. Concordant Patterns of Genome-Wide DE with TCGA

We next examined the concordance of ranked gene lists derived from each MMD based on the degree of DE with matching TCGA cohorts. As all MMDs were assayed with the same Affymetrix platform and processed with uniform R framework, same number of genes (20,456 genes) were present in the final ranked gene lists. To dispel any bias that could be introduced from different number of genes probed with various profiling technologies, we compared only the ranked gene lists that were common for both MMD and TCGA (Table 2).

**Table 2.** Number of genes included in the final ranked gene list in MMD and TCGA for cross-platform RRHO analysis.

| MMD | Gene *N* | TCGA | Gene *N* | Commone gene *N* |
| --- | --- | --- | --- | --- |
| Lung | 20546 | LUAD | 15404 | 14101 |
| Pancreas | 20546 | PAAD | 15533 | 14239 |
| Prostate | 20546 | PRAD | 14976 | 13720 |
| Kidney | 20546 | KICH | 15064 | 13802 |
| Stomach | 20546 | STAD | 15418 | 14104 |
| Colon | 20546 | COAD | 14785 | 13571 |
| Breast | 20546 | BRCA | 15165 | 13887 |
| Liver | 20546 | LIHC | 14138 | 13022 |
| Bladder | 20546 | BLCA | 15033 | 13771 |

A rank-rank hypergeometric overlap (RRHO) algorithm [20] was used to identify and visualize statistically significant overlap between signatures of all protein-coding genes (Figure 3). This threshold-free approach allows the analysis of genes with low expression that would be missed by other conventional methods which depend on a user-defined, fixed cutoff. Each ranked gene list was loaded into a web-based executable version of RRHO (http://systems.crump.ucla.edu/rankrank/rankranksimple.php) to generate Benjamin-Yekutieli corrected hypergeometric matrix with step size of 300.

**Figure 3.**
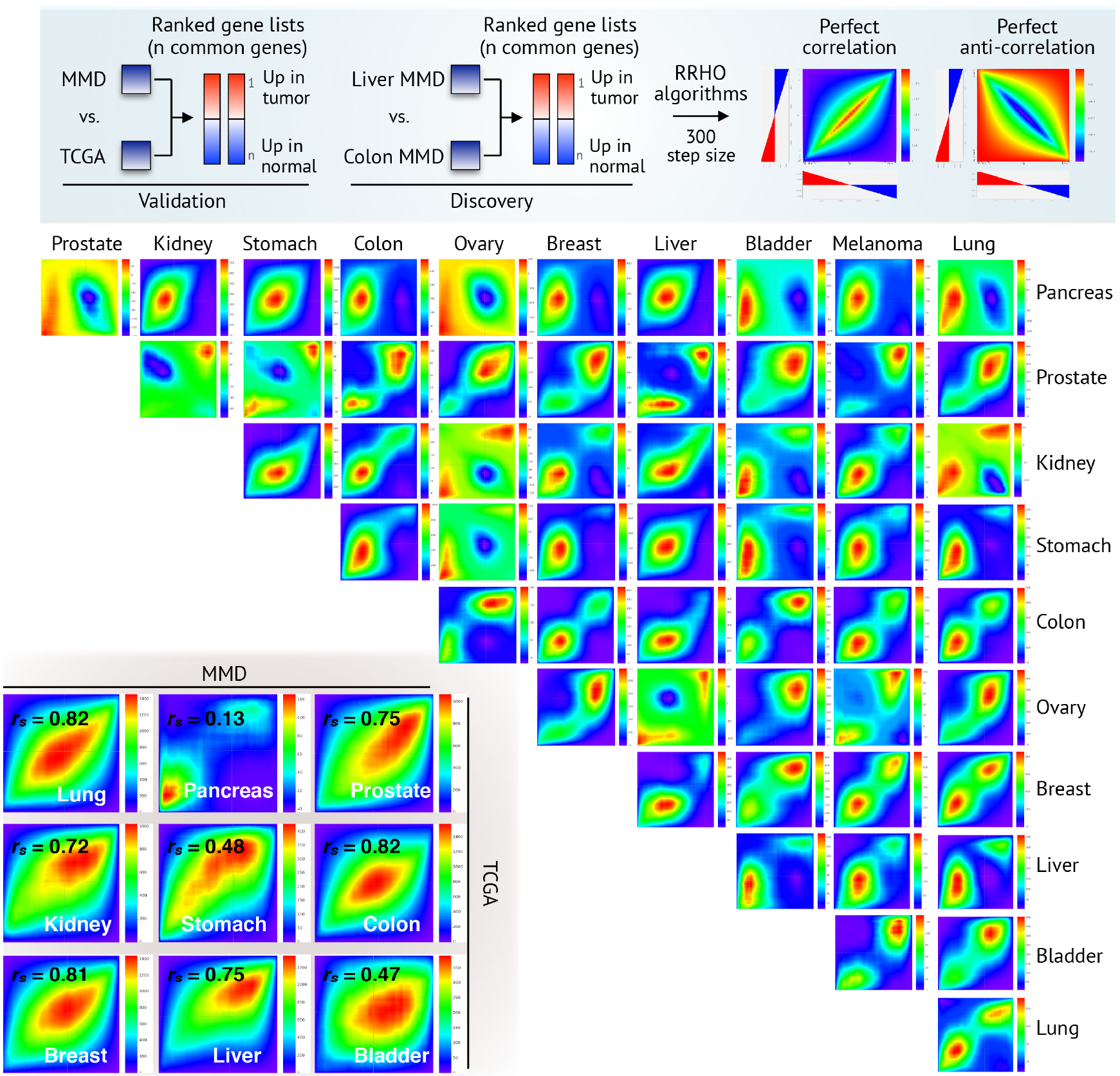
Concordant patterns of genome-wide differential expression with TCGA. Rank-rank hypergeometric overlap (RRHO) heatmaps are drawn to visualise the cross-platform concordance between microarray (MMD) and RNA-seq (TCGA) in deriving ranked gene lists (bottom left). All protein-coding genes available in the preprocessed data were ranked by the degree of differential expression computed in respective MMDs and TCGA cohorts (cancer versus normal). MMD-derived RRHO heatmaps are also shown to demonstrate potential use of MMDs for cross-cancer analysis. *r*_*s*_ = Spearman’s correlation coefficient (Spearman’s correlation *P*-value < 0.001).

A significant overlap with TCGA-derived ranked lists was observed in lung, prostate, kidney, colon, breast, and liver MMD (Spearman’s correlation coefficient, *r*_*s*_, ranging from 0.72 to 0.82). The weak correlation observed from pancreas, stomach, and bladder MMD were likely due to a relatively small number of tumor-free tissues available in respective TCGA datasets (Table 1). These two additional quality control metrics implemented in the integrated R pipeline served to ensure the validity and reproducibility of DE measures of MMDs. Additionally, we generated RRHO heatmaps to demonstrate potential use of MMDs in evaluating the extent to which transcriptomic signatures are shared across cancer type; Spearman’s correlation coefficients across 11 cancer types are shown in Table 3. Uniform preprocessing techniques used to build MMDs further enable cross-cancer analysis. It should be of particular interest to those developing scoring metrics or index based on certain molecular features for any given sample to establish spectrum across various carcinomas, revealing inter-tumor variation, or heterogeneity.

**Table 3.** Cross-cancer genome-wide DE analysis using MMD: Spearman’s correlation coefficients across 11 cancer types

| MMD | Lung | Pancreas | Prostate | Kidney | Stomach | Colon | Ovary | Breast | Liver | Bladder | Melanoma |
| --- | --- | --- | --- | --- | --- | --- | --- | --- | --- | --- | --- |
| Lung | 1.000 | -0.002 | 0.311 | -0.027 | 0.161 | 0.315 | 0.397 | 0.482 | 0.250 | 0.300 | 0.150 |
| Pancreas | -0.002 | 1.000 | -0.104 | 0.362 | 0.542 | 0.176 | -0.191 | 0.159 | 0.360 | 0.030 | 0.165 |
| Prostate | 0.311 | -0.104 | 1.000 | -0.044 | 0.001 | 0.139 | 0.231 | 0.340 | 0.072 | 0.349 | 0.077 |
| Kidney | -0.027 | 0.362 | -0.044 | 1.000 | 0.362 | 0.188 | -0.053 | 0.107 | 0.270 | 0.087 | 0.219 |
| Stomach | 0.161 | 0.542 | 0.001 | 0.362 | 1.000 | 0.439 | -0.011 | 0.229 | 0.396 | 0.105 | 0.280 |
| Colon | 0.315 | 0.176 | 0.139 | 0.188 | 0.439 | 1.000 | 0.145 | 0.250 | 0.309 | 0.158 | 0.238 |
| Ovary | 0.397 | -0.191 | 0.231 | -0.053 | -0.011 | 0.145 | 1.000 | 0.348 | 0.003 | 0.287 | 0.072 |
| Breast | 0.482 | 0.159 | 0.340 | 0.107 | 0.229 | 0.250 | 0.348 | 1.000 | 0.261 | 0.357 | 0.289 |
| Liver | 0.250 | 0.360 | 0.072 | 0.270 | 0.396 | 0.309 | 0.003 | 0.261 | 1.000 | 0.108 | 0.234 |
| Bladder | 0.300 | 0.030 | 0.349 | 0.087 | 0.105 | 0.158 | 0.287 | 0.357 | 0.108 | 1.000 | 0.136 |
| Melanoma | 0.150 | 0.165 | 0.077 | 0.219 | 0.280 | 0.238 | 0.072 | 0.289 | 0.234 | 0.136 | 1.000 |

## MMD-based Machine Learning for Predictive Medicine

### 1. Cancer Classifier

Publicly-accessible data repositories, such as GTEx [21], TCGA [22], HPA [23], and ArrayExpress [24], host genome-wide expression profiles assayed with various profiling technologies. Having sufficient read depth [9], higher resolution [10], higher dynamic range [11], and lower technical variation [25], RNA-seq is increasingly the platform of choice in translational-biomarker studies. Paralleling this trend, crossplatform normalization tools to facilitate comparison of data from different platforms continue to be developed. The techniques specifically designed to transform RNA-seq data to make it compatible with microarray data include PREBS [26], VOOM [27], and TDM [11]. Other conventional methods also exist in dealing with such ‘dataset shifts’ [28], such as quantile normalization, log_2_ transformation, and nonparanormal transformation [11]. It is such algorithms – not just datasets – that will prove useful when combined with machine learning.

To demonstrate potential use of MMD for machine learning applications, we developed a new cancer classifier by training each cancer type-specific MMD and tested on respective RNA-seq-acquired TCGA using supervised machine learning (Figure 4a). Using the *glmnet* package [29], we performed LASSO multinomial logistic regression [30] with 100 fold cross-validation (CV) to build best predictive model in identifying cancer from normal samples. By comparing a simple scaling, log_2_, and TDM transformation methods, we first found that TDM transformation best fits the reference MMD data distribution (Figure 4b). Predictive model built from each MMD was then tested directly on TDM-transformed-TCGA dataset. Except for breast MMD, all MMDs achieve an average accuracy of 0.96 in classifying TCGA cancers (ranging from 0.913 to 0.997; Figure 4c).

**Figure 4.**
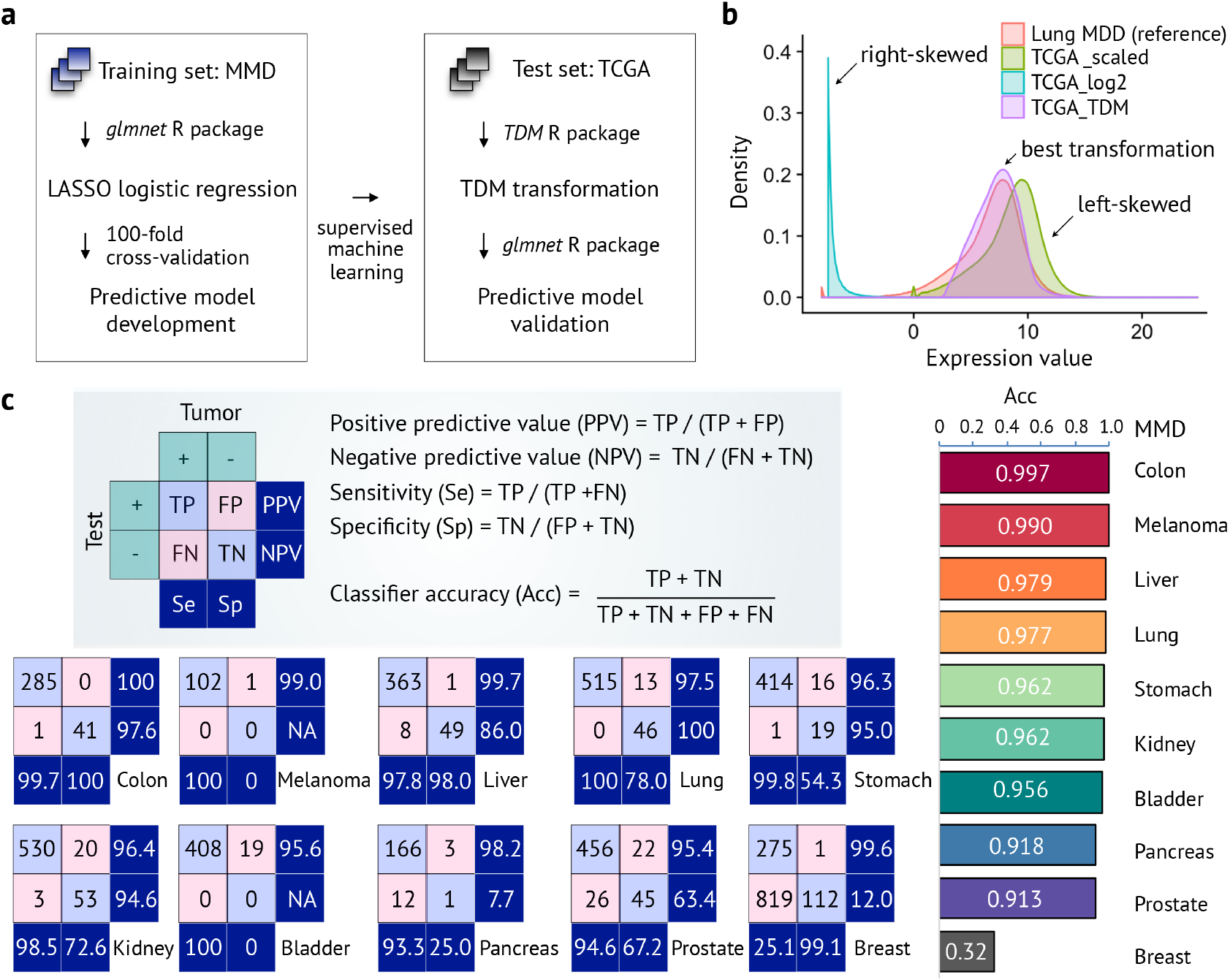
Supervised machine learning classifies cancer. (a) Schematic workflow: cancer classifier is built and trained from each microarray-acquired-MMD and tested on matching RNA-seq-acquired TCGA using LASSO logistic regression. (b) TDM-transformed testing data (TCGA LUAD) best fits the training data distribution (lung MMD). (c) Except breast MMD, generated MMDs achieve an average accuracy of 0.96 in classifying cancer from normal tissues.

Such high classifier accuracy achieved by our resource of large-scale microarray data suggests that machine learning derived models built from MMD can be applied directly to existing RNA-seq-acquired data or prospectively generated data for clinical endpoint prediction. Given that the performance of machine learning algorithms is highly dependent on the quantity and quality of input data, requiring considerable number of observations [31], MMD will better help us to develop thousands of rich predictive indicators for diagnosis and prognosis.

### 2. Pan-cancer Immunogenomic Analyses

Remarkable clinical success in immunotherapy has changed the way we treat cancer. At the same time, the majority of patients do not respond to immunotherapy. This has raised the question of what drives such heterogeneity in sensitivity across various carcinomas. Our data source focusing on multiple datasets will enable the direct comparisons to build such predictive models for immunotherapy. TCGA data, for example, are increasingly mined to find prognostic influence of estimated composition immune cell infiltrates [32, 33], predictors of checkpoint blockade response [34], and association of neoantigens with survival [35] or with estimated cytolytic activity [36]. Computational tools for large-scale immunogenomics further facilitate high-resolution analyses of the tumor-immune interface (see [37] for summary of existing analytical pipelines). Thus, we further tested if our curated databases could provide the basis and make use of such computational infrastructure.

Applying CIBERSORT (http://cibersort.standford.edu/) [12], we first computed the estimated fraction of 22 immune cell populations using the beta version (Figure 5a). Only tumor samples were extracted for the deconvolution in each case. Having whole-transcriptome profiles of over 1,500 samples, breast, colon, and lung MMD exceeded the maximum load file capacity (500 MB); 1,000 tumor samples were thus randomly selected to generate the input “mixture” file. Each MMD was ran with 100 permutations for LM22 (22 immune cell types) signature gene. Using the relative abundance of immune cell infiltrate frequencies, we further estimated the functional activity of six distinct cell types, for their potential role in specific killing of tumor cells upon immunotherapy [5].

**Figure 5.**
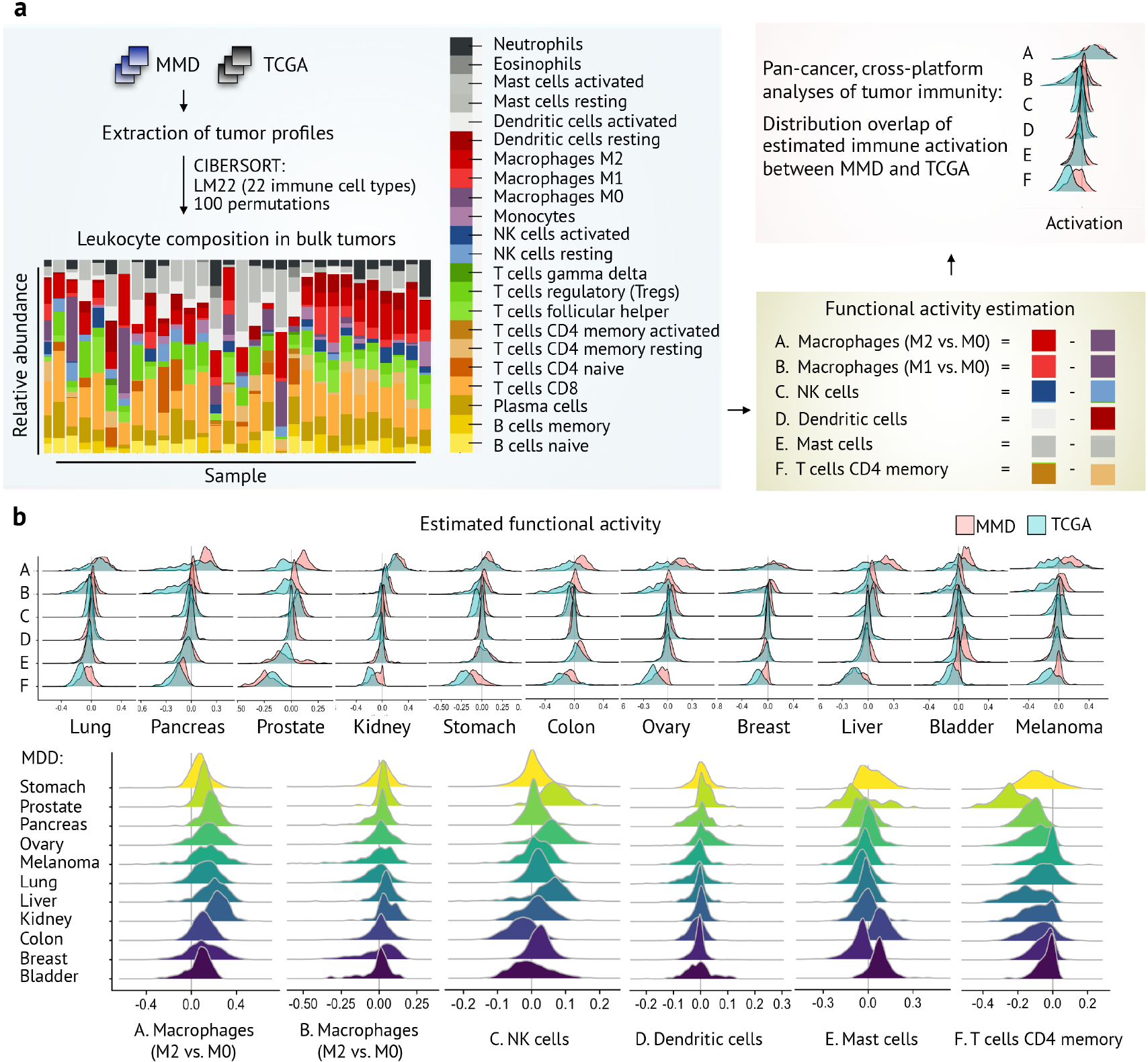
MMD reveals the immune landscape in human cancer. (a) Schematic workflow: MMD provides the basis to dissect the tumor-immune interface. Relative abundance of 22 immune cell types and functional activation of six distinct cell types of immune infiltrates were estimated using CIBERSORT. (b) Highly overlapping immune landscape in MMD-derived bulk tumors with respective TCGA cohorts (top). MMD allows cross-cancer evaluation of tumor immunity across 11 cancer types, given the derivation from a uniform R processing framework (bottom).

To validate cross platform, we repeated this procedure and examined the degree of distribution overlap with matching TCGA cohorts. Having highly overlapping immune landscape with TCGA across all cancer types, MMD further facilitates cross-cancer analysis of specific cellular composition of immune infiltrates (Figure 5b). There are also several quantitative measures, or indices, of genomic variation associated with disease progression (e.g., stemness [5], EMT [38], and ECM [1]) that can be applied to large-scale MMD data with deconvolution method.

To demonstrate its applications, we examined the correlation of matrisome pattern that accompanies tumor progression with the composition of immune infiltrates in over 30,000 samples using both MMD and TCGA. Regardless of cancer type, significant enrichment of certain immune cell types was found in tumors with deregulated matrisome, possibly suggesting potential role of ECM molecules in shaping immunosuppressive microenvironment (unpublished). As such, multi-platform, pan-cancer immunogenomic analyses of the present MMD data may reveal the interconnectivity of such distinct phenotypes and explain how tumors shape protumoral and immunosuppressive microenvironment to promote their growth and immune escape.

## Conclusion

Our open resource of MMDs combined with machine learning empowers precision cancer medicine by facilitating

1. pan-cancer analysis of transcriptome, allowing quantification of phenotypic spectrum of interest across various carcinomas,
2. multi-platform assessment of biomarker-derived predictors with existing or new large-scale data acquired from different profiling technologies,
3. development of new clinical predictive model and validation with (or application to) independent validation cohort using machine learning approach, and
4. optimization of machine learning and cross-platform normalization algorithms requiring large-scale training and test data.

## Abbreviations

PCA: principal component analysis
DE: differential expression
LUAD: lung adenocarcinoma
PAAD: pancreatic adenocarcinoma
PRAD: prostate adenocarcinoma
KIRC: kidney renal clear cell carcinoma
STAD: stomach adenocarcinoma
COAD: colon adenocarcinoma
OV: ovarian serous cystadenocarcinoma
BRCA: breast carcinoma
LIHC: liver hepatocellular carcinoma
BLCA: bladder carcinoma
SKCM: skin cutaneous melanoma
TCGA: the Cancer Genome Atlas
GTEx: Genotype-Tissue Expression
HPA: the Human Protein Atlas
GEO: Gene Expression Omnibus
NCBI: National Center for Biotechnology Information
RRHO: rank-rank hypergeometric overlap

## Declarations

## Acknowledgements

This work was conceived and carried out at the MechanoBioEngineering laboratory at the Department of Biomedical Engineering, National University of Singapore (NUS). We acknowledge support provided by the National Research Foundation, Prime Minister’s Office, Singapore under its Research Centre for Excellence, and Mechanobiology Institute at NUS. S.B.L. acknowledges scholarship and support from NUS Graduate School for Integrative Sciences and Engineering (NGS).

## Availability of Supporting Data

All processed whole transcripome profiles of 11 MMDs (with associated clinical metadata) have been deposited at ArrayExpress under accession codes E-MTAB-6690 (pancreatic cancer), E-MTAB-6691 (ovarian cancer), E-MTAB-6692 (renal cancer), E-MTAB-6693 (gastric cancer), E-MTAB-6694 (prostate cancer), E-MTAB-6695 (liver cancer), E-MTAB-6696 (bladder cancer), E-MTAB-6697 (melanoma cancer), E-MTAB-6698 (colorectal cancer), E-MTAB-6699 (lung cancer), and E-MTAB-6703 (breast cancer).

## Competing Interests

The authors declare no competing interests.

## Author Contributions

S.B.L., S.J.T., W.-T.L. and C.T.L. conceptualized and designed the study. S.B.L. developed the R pipeline to generate MMDs. S.B.L., S.J.T., W.-T.L and C.T.L. analyzed and interpreted the data. S.B.L., S.J.T., W.-T.L and C.T.L. reviewed and contributed to the manuscript.

